## Supplemental Figure S1 for "DGRPool: A web tool leveraging harmonized *Drosophila* Genetic Reference Panel phenotyping data for the study of complex traits"

### Editing study

PAPER INFORMATION

Context-dependent genetic architecture of Drosophila life span

*[10.1371/journal.pbio.3000645][FBrf0245198]*

**Title**

Context-dependent genetic architecture of Drosophila life span

**Authors**

[{"initials":"W","fname":"Wen","lname":"Huang","aff_ids":[]},{"initials":"T","fname":"Terry","lname":"Campbell","aff_ids":[]},{"initials":"M A","fname":"Mary Anna","lname":"Carbone","aff_ids":[]},{"initials":"W. E","fname":"W. Elizabeth","lname":"Jones","aff_ids":[]},{"initials":"D","fname":"Desiree","lname":"Unselt","aff_ids":[]},{"initials":"R R. H.","fname":"Robert R. H.","lname":"Anholt","aff_ids":[]},{"initials":"T F. C.","fname":"Trudy F. C.","lname":"MacKay","aff_ids":[]}]

CURATION PROCESS

#### Status

Validated

#### Comments from curator

Phenotyping data are available from the <a href="https://doi.org/10.1371/journal.pbio.3000645.s001">S1 table</a> found in the <a href="https://journals.plos.org/plosbiology/article?id=10.1371/journal.pbio.3000645#sec019">Supporting information</a> section. <b>Of note</b>, raw data is also available from <a href="https://github.com/qgg-lab/dgrp-lifespan/">GitHub</a>, so we primarily uploaded this data, instead of the summarized data.<br/><br/>

The authors also created an Advanced Intercross Population (AIP) from a subset of extreme long-living DGRP lines that were maintained for over 100 generations with a large effective population size before performing "extreme quantitative trait locus (xQTL)" mapping for life span.

ABSTRACT

#### Description

In this study, the authors quantified variation in life span in males and females reared in 3 thermal environments. Quantitative genetic analyses of life span and the micro-environmental variance of life span in the DGRP revealed significant genetic variance for both traits within each sex and environment, as well as significant genotype-by-sex interaction (GSI) and genotype-by-environment interaction (GEI). We only kept the micro-environmental variance (ln<i>σ</i><sub>ε</sub>) summary calculation from the authors, but it's a simple logarithmic transformation of the standard deviation.

ANNOTATION

#### Other external data available

PRJNA577841: Whole Genome Sequencing of whole flies (24 samples) taken from Advanced Intercross Population (AIP) from a subset of DGRP lines with longest living (M & F, 3 different temperature environments)

#### Categories

Select a category to add it

Evolution ✕ Life history traits ✕ Aging ✕

FILTERING

#### Deactivate phenotypes

Select a phenotype to deactivate

Block ✕

#### Deactivate DGRP lines

Select a DGRP line to deactivate
