## Supplementary figures and images for "DGRPool: A web tool leveraging harmonized *Drosophila* Genetic Reference Panel phenotyping data for the study of complex traits"

### Supplemental Figure S2

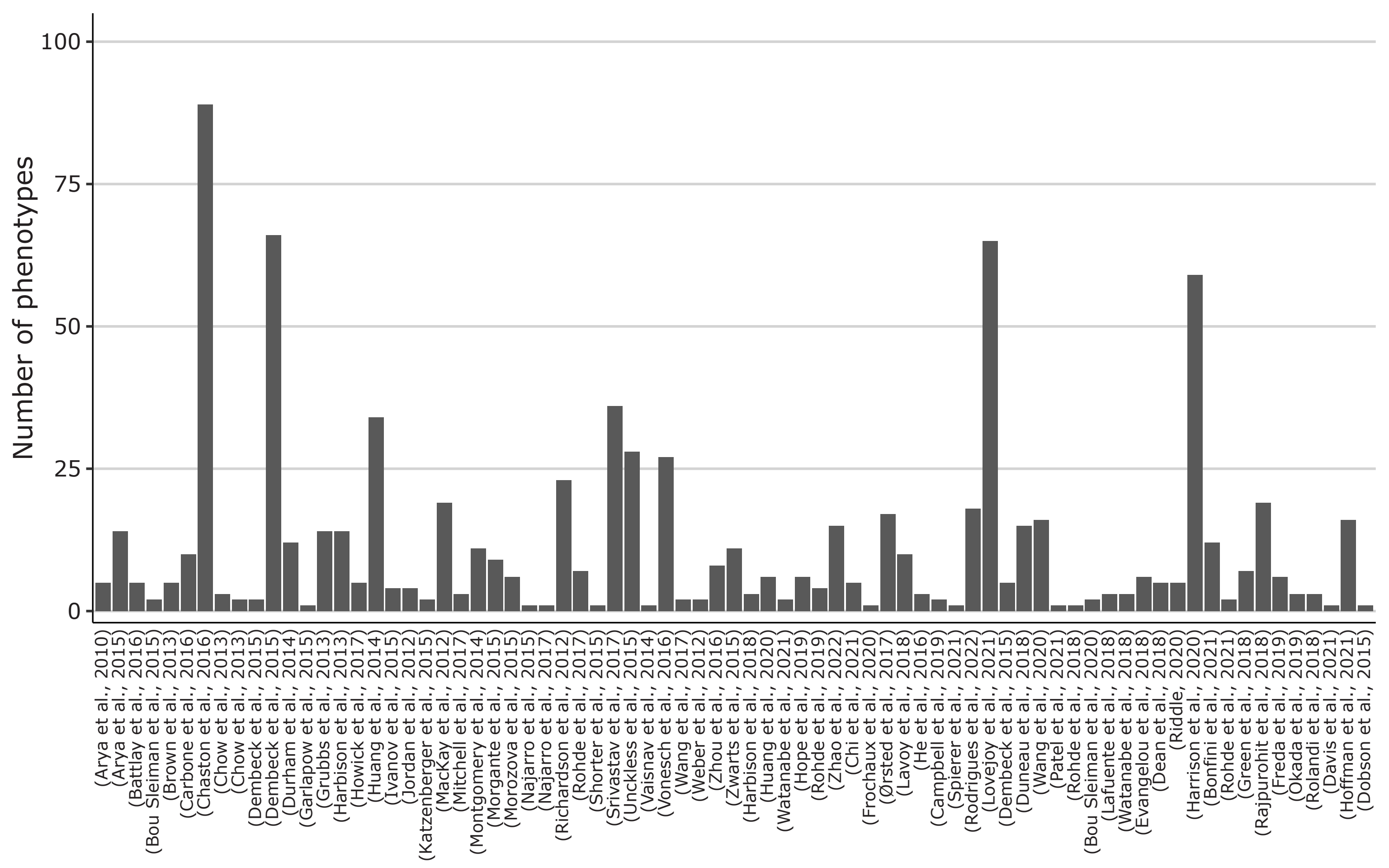

### Supplemental Figure S3

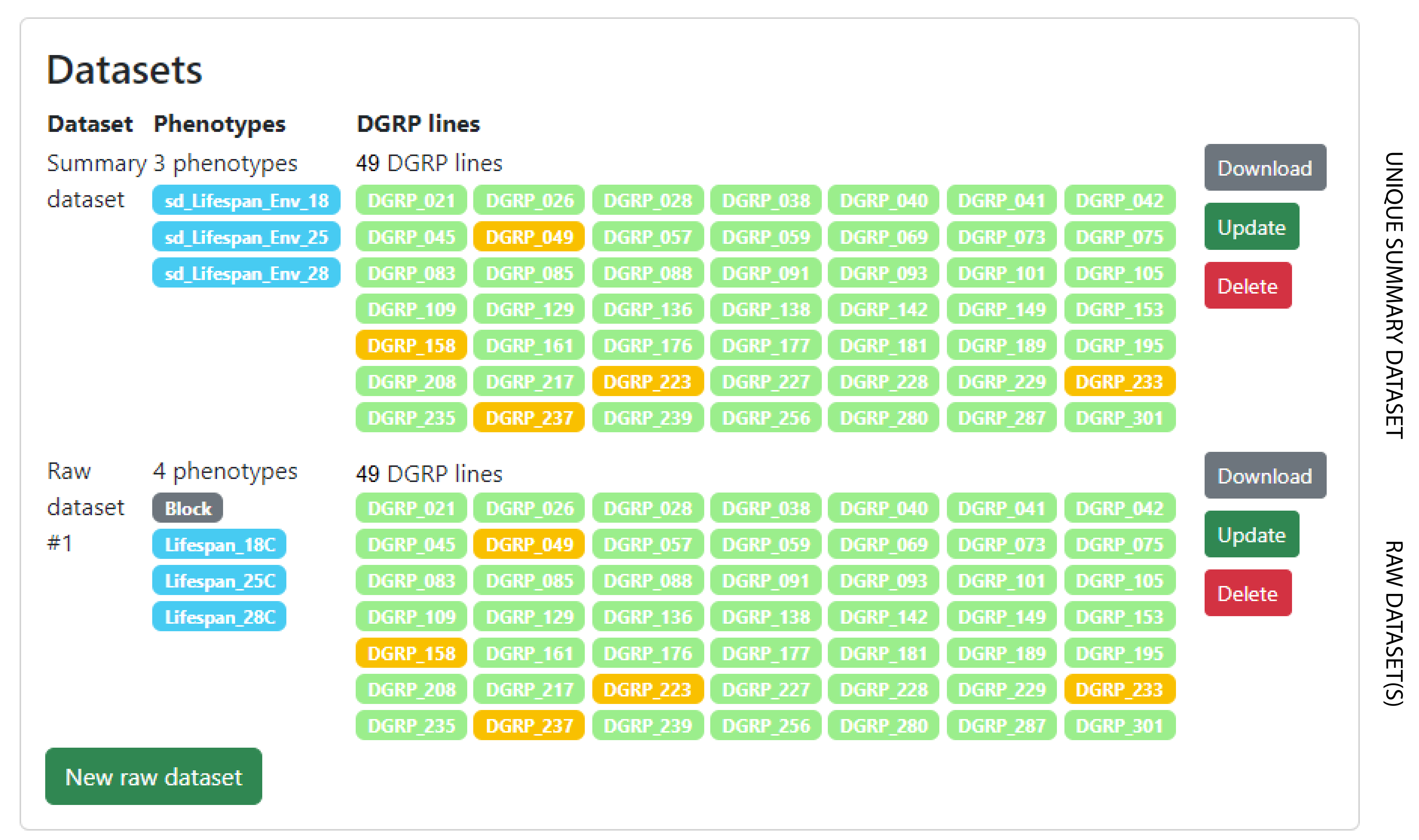

### Supplemental Figure S5

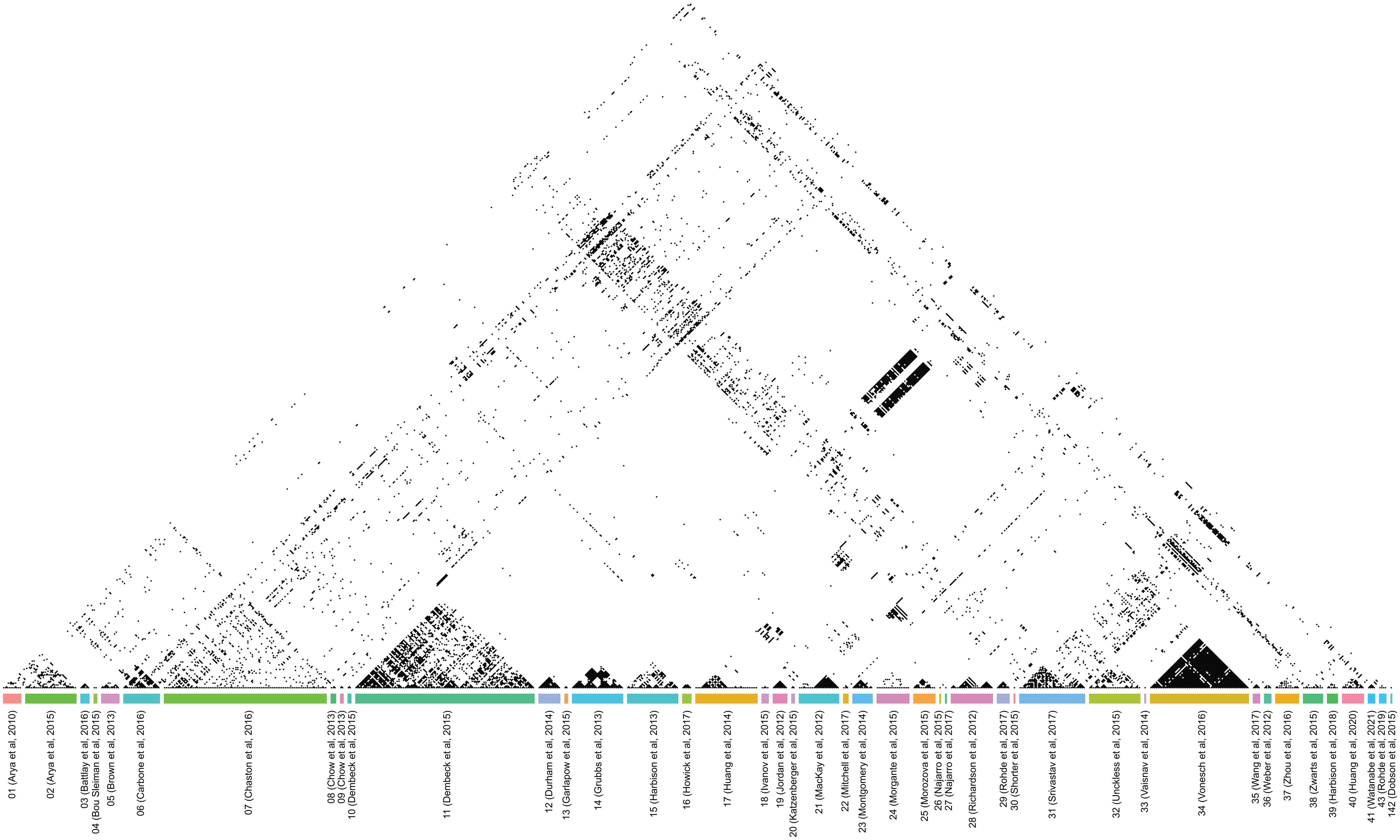

### Supplemental Figure S6

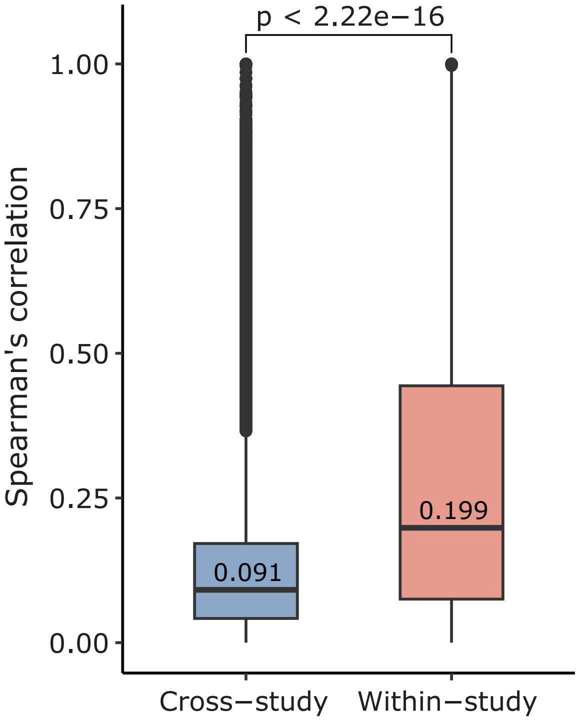

### Supplemental Figure S7

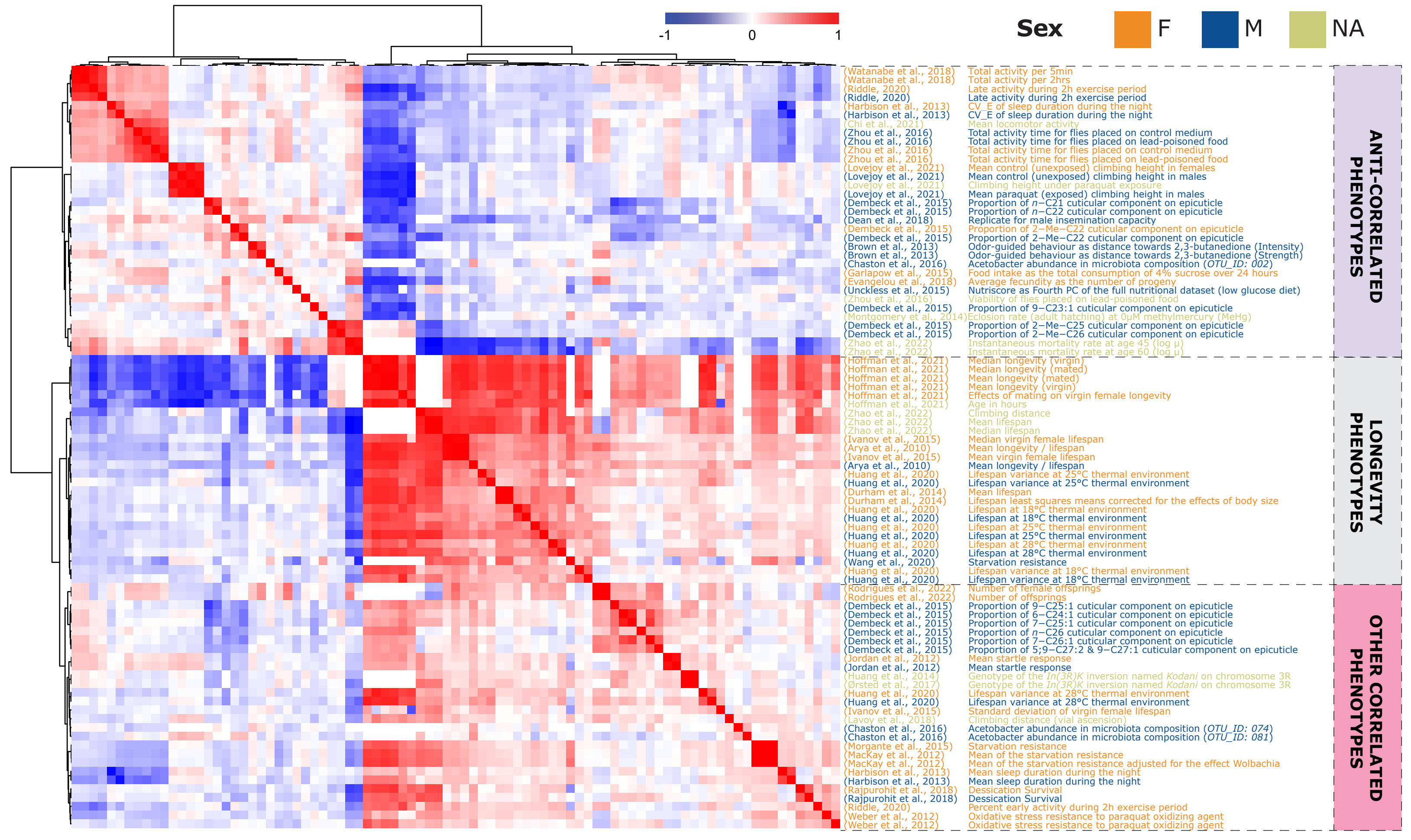

### Supplemental Figure S9

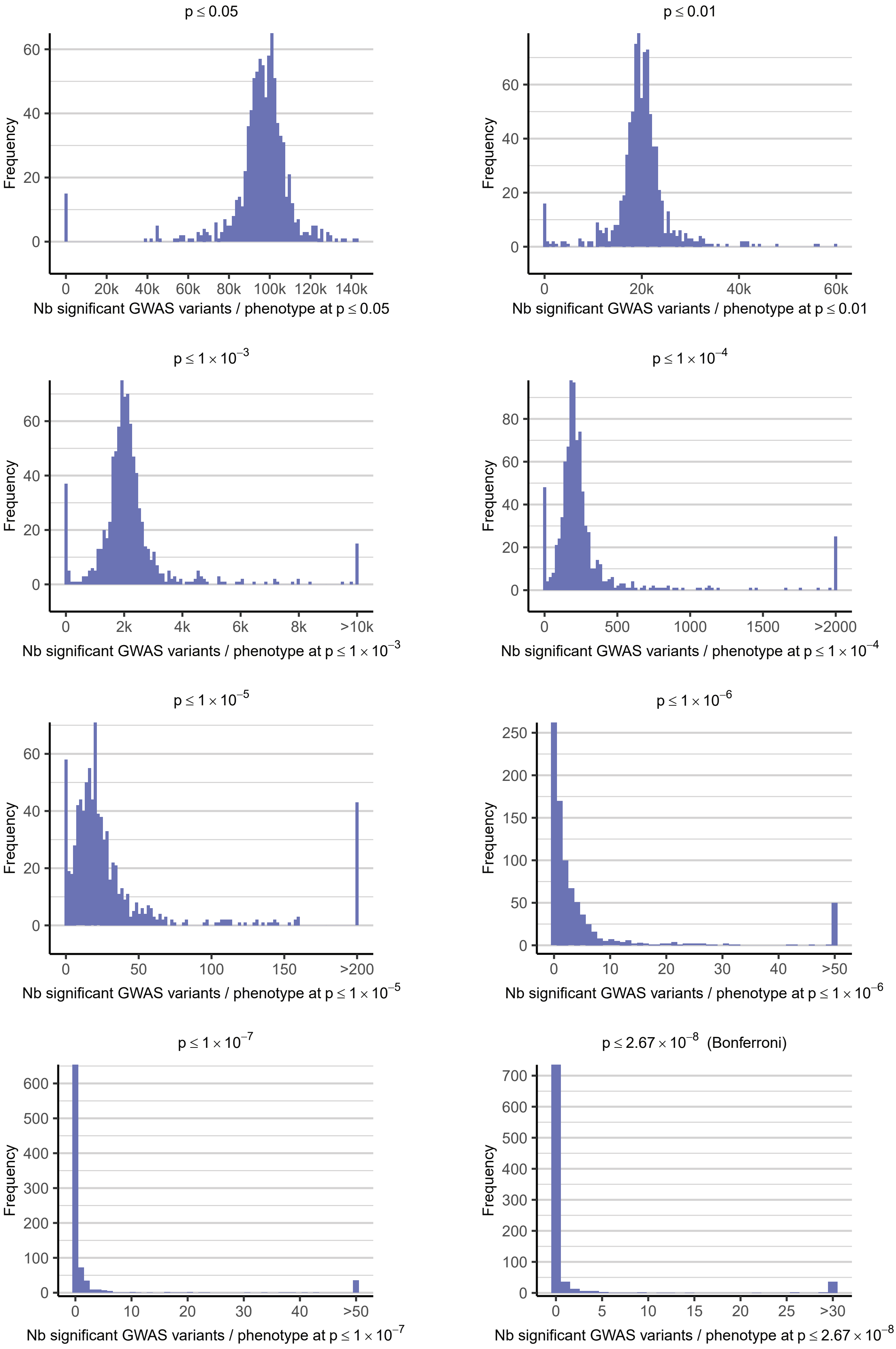
