## Supplemental Figure S4 for "DGRPool: A web tool leveraging harmonized *Drosophila* Genetic Reference Panel phenotyping data for the study of complex traits"

For robustness of the results we restrict by default the available phenotypes to curated studies but if you prefer to be able to access the whole dataset that was automatically extracted, you can change this behaviour below.

☑ Only curated studies

Female 657 | Male 657

### Results for mn_Longevity Female

Show 25 entries Search:

| Study | phenotype | sex | Cor. | p | FDR | DGRP overlap |
|---|---|---|---|---|---|---|
| Arya et al., 2010 | mn_Longevity | female | 1 | 0 | 0 | 218 |
| Ivanov et al., 2015 | mn_Lifespan | female | 1 | 0 | 0 | 125 |
| Ivanov et al., 2015 | md_Lifespan | female | 0.965 | $1.584e^{-54}$ | $4.757e^{-52}$ | 93 |
| Arya et al., 2010 | mn_Longevity | male | 0.602 | $6.363e^{-23}$ | $1.433e^{-20}$ | 218 |
| Huang et al., 2020 | Lifespan_25C | male | 0.553 | $3.467e^{-11}$ | $6.248e^{-9}$ | 123 |
| Huang et al., 2020 | sd_Lifespan_Env_25 | female | 0.532 | $2.523e^{-10}$ | $3.789e^{-8}$ | 123 |
| Huang et al., 2020 | Lifespan_25C | female | 0.496 | $5.482e^{-9}$ | $7.056e^{-7}$ | 123 |
| Huang et al., 2020 | sd_Lifespan_Env_25 | male | 0.49 | $8.762e^{-9}$ | $9.868e^{-7}$ | 123 |
| Huang et al., 2020 | sd_Lifespan_Env_18 | female | 0.457 | $1.555e^{-7}$ | $1.557e^{-5}$ | 120 |
| Huang et al., 2020 | sd_Lifespan_Env_18 | male | 0.438 | $5.649e^{-7}$ | $5.090e^{-5}$ | 120 |
| Huang et al., 2020 | Lifespan_18C | female | 0.414 | $2.610e^{-6}$ | $2.138e^{-4}$ | 120 |
| Huang et al., 2020 | Lifespan_18C | male | 0.411 | $3.030e^{-6}$ | $2.275e^{-4}$ | 120 |
| Huang et al., 2020 | Lifespan_28C | male | 0.41 | $5.435e^{-6}$ | $3.767e^{-4}$ | 115 |
| Zhou et al., 2016 | mn_Tot_Activity_Lead | female | -0.417 | $1.169e^{-5}$ | $7.524e^{-4}$ | 103 |
| Durham et al., 2014 | mn_Lifespan | female | 0.351 | $6.270e^{-5}$ | 0.004 | 124 |
| Durham et al., 2014 | lsm_Lifespan | female | 0.349 | $7.080e^{-5}$ | 0.004 | 124 |
| MacKay et al., 2012 | mn_StarvationRes_ADJ | female | 0.357 | $1.025e^{-4}$ | 0.005 | 113 |
| Morgante et al., 2015 | StarvationRes | female | 0.332 | $1.293e^{-4}$ | 0.006 | 128 |
| MacKay et al., 2012 | mn_StarvationRes | female | 0.346 | $1.753e^{-4}$ | 0.008 | 113 |
| Dembeck et al., 2015 | Cuticul_9_C25_1 | male | 0.346 | $4.830e^{-4}$ | 0.022 | 98 |
| Zhou et al., 2016 | mn_Tot_Activity_Lead | male | -0.332 | $6.125e^{-4}$ | 0.026 | 103 |
| Huang et al., 2020 | sd_Lifespan_Env_28 | female | 0.313 | $6.526e^{-4}$ | 0.027 | 115 |
| Dembeck et al., 2015 | Cuticul_2_Me_C26 | male | -0.335 | $7.391e^{-4}$ | 0.028 | 98 |
| Huang et al., 2020 | Lifespan_28C | female | 0.307 | $8.386e^{-4}$ | 0.03 | 115 |
| Huang et al., 2020 | sd_Lifespan_Env_28 | male | 0.304 | $9.606e^{-4}$ | 0.033 | 115 |

Showing 1 to 25 of 657 entries
