## Supplemental Figure S8 for "DGRPool: A web tool leveraging harmonized *Drosophila* Genetic Reference Panel phenotyping data for the study of complex traits"

### mn_Surv_Azinpho_0_25 phenotype

Back to phenotype

GWAS results were precomputed for all phenotypes using PLINK2 [v2.00a3LM (1 Jul 2021)] and corrected for the 6 main covariates defined in Huang et al., 2014. These covariates are the same than the ones used in the DGRP2 website.

Unclassified sex 2222

Filters 0

GWAS results were computed on dm3 genome.

If you want to make an independent analysis with DGRP2 or PLINK2, you can find here prepared input files for this phenotype: DGRP2 results PLINK2 results

#### Phenotype distribution and association to known covariates

Full results

**Shapiro-Wilk test of normality** (p-value=2.855e$^{-12}$)
Null Hypothesis is rejected (p<=0.05).
Interpretation: NOT NORMAL distribution of this phenotype

We tested for association of this phenotype with 6 known covariates Huang et al., 2014 using both ANOVA and Kruskal-Wallis association tests.

**ANOVA test**

| Covariate | p-value | Significance |
|---|---|---|
| In_2R_NS | 0.001511 | ★★★☆ |

**Kruskal-Wallis test**

| Covariate | $\chi^2$ | p-value | Significance |
|---|---|---|---|
| In_2L_t | 5.1403 | 0.0765 | ★☆☆☆ |
| In_2R_NS | 8.2249 | 0.0164 | ★★☆☆ |

#### QQplot

PNG PDF

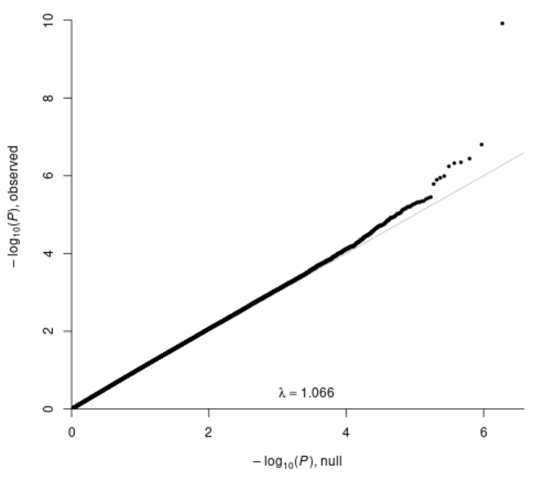


#### Manhattan plot

PNG PDF

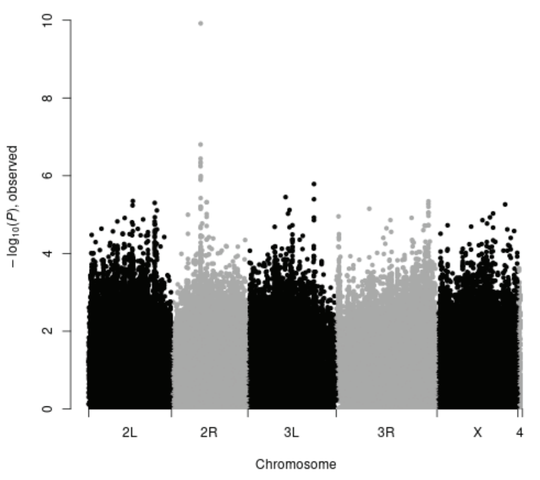


#### GWAS results

GWAS results

2222 significant hits at p-value < 0.001. The 1000 top results are displayed in the table below.

We performed a gene enrichment calculation (Fisher's exact test) of all GWAS-associated genes at p < 0.001. Genesets were taken from two database: the Gene Ontology (GO) and the FlyBase phenotypes. GO enrichment FlyBase phenotype enrichment

**Variant impact** HIGH MODERATE LOW MODIFIER
Show 100 entries

Search:

| Chromosome | Position | ID | Reference | Variant | p-value | FDR_BH | Transcript annotation | Binding site annotation | |
|---|---|---|---|---|---|---|---|---|---|
| 2R | 8072884 | 2R_8072884_INS [dm3] 2R_12185379_INS [dm6] | G | GAGTTACGGGTGCCTCCGTT... | 1.204e$^{-10}$ | 2.259e$^{-4}$ | UTR_3_PRIME CG13175 & 1 other; UPSTREAM Cyp6g1 | regulatory region REDfly_CRMs | Boxplot PheWAS |
| 2R | 8032968 | 2R_8032968_DEL [dm3] 2R_12145463_DEL [dm6] | AA | A | 1.576e$^{-7}$ | 0.148 | UPSTREAM CG13178 & 1 other; DOWNSTREAM CG8878; INTERGENIC | | Boxplot PheWAS |
